# Bacterial phylogenetic markers in lake sediments as evidence for historical hemp retting

**DOI:** 10.1101/2022.08.03.502709

**Authors:** Valentí Rull, Oriol Sacristán-Soriano, Alexandre Sànchez-Melsió, Carles M. Borrego, Teresa Vegas-Vilarrúbia

## Abstract

Documenting prehistoric and historical hemp retting for fiber extraction is important in the study of human uses of this iconic plant and its cultural implications. In paleoecology, hemp retting is usually inferred from indirect proxies, notably anomalously high percentages of *Cannabis* pollen in lake sediments, but some recent studies have also used specific molecular biomarkers (cannabinol, *Cannabis* DNA) as a more straightforward evidence. Here we provide direct evidence of hemp retting by identifying phylogenetic signatures (16S *rRNA* genes) from pectinolytic bacteria actually responsible for the fermentation process that separates the fiber from the stalk, namely *Bacillus, Clostridium, Escherichia, Massilia, Methylobacterium, Pseusomonas, Rhizobium* and *Rhodobacter*. These analyses have been performed in the sediments from an Iberian lake previously considered as an important hemp retting site during the last five centuries, on the basis of *Cannabis* pollen abundances. The good match between biomarker and pollen evidence, in the context of the recent historical development of hemp industry in Spain, can be useful to interpret paleoecological records from other similar lakes in the way toward a more regional view on the introduction, spreading, uses and associated cultural connotations of *Cannabis* in the Iberian Peninsula within European and Mediterranean contexts.

## Introduction

*Cannabis* is one of the most ancient crops and has been an integral part of human life since its early Neolithic domestication. Human-mediated selection of this plant has created a wide array of varieties and biotypes for diverse uses, including hemp fiber, human and animal feed, biofuel, medicine, recreational drugs and ritual activities, among others (Clarke & Merlin 2013). An updated review on the evolutionary origin, early expansion, domestication and anthropogenic diffusion of *Cannabis* is available at Rull (2022). The fossil record of *Cannabis* is largely based on pollen, but the morphological similarity between *Cannabis* and *Humulus* pollen has been a handicap for the correct identification of these Cannabaceae genera in past records. As a result, different authors refer to the fossil representatives of this pollen type using unspecific names such as *Cannabis*/*Humulus, Cannabis-type, Humulus*-type or Cannabaceae. Several morphological traits have been proposed to distinguish *Cannabis* and *Humulus* pollen but there is no universal agreement in this respect (McPartland et al., 2018).

The finding of unexpectedly large amounts of *Cannabis*/*Humulus* pollen is not uncommon in lake sediments corresponding to the last millennia, which is usually attributed to hemp retting, a fermentation process needed to separate the epidermal hemp fibers from the stems by degrading cementing materials, especially pectin (Akin, 2013). Currently, hemp retting is performed industrially with specific bacterial formulations but in historical times, fiber extraction was accomplished by immersion of hemp plants in lakes and ponds, where pectin was naturally degraded by the autochthonous pectinolytic bacterial flora. In this way, the pollen of male hemp plants was directly incorporated into aquatic sediments and was overrepresented in the resulting fossil assemblages. *Cannabis*/*Humulus* pollen values around 80-90% are not unusual in these cases, but abundances of 15-25% have been considered enough for interpreting hemp retting (review in McPartland et al., 2018). However, pollen abundance solely is an indirect evidence for hemp retting because the high dispersion capacity of this anemophilous pollen favors its occurrence in significant percentages on sediments from lakes not used for hemp retting (Rull et al., 2017). Hemp fiber counting has also been used as a proxy for hemp retting intensity but discrepancies with pollen records and potential interferences caused by local hemp cultivation have been identified (Iwańska et al., 2022).

Recently, paleoecological applications of biomolecular methods have provided more straightforward evidence for the identification of hemp retting in past lake sequences. For example, Lavrieux et al. (2013) used sedimentary cannabinol, a specific *Cannabis* compound absent in pollen grains, as a reliable indicator of retting. Similarly, Giguet-Covex et al. (2019) utilized sedimentary DNA from *Cannabis sativa* to trace the history of hemp retting. In both cases, comparisons between specific biomarkers and pollen helped not only to reconstruct the retting history of the studied sites but also provided tools to reliably discern between *Cannabis* and *Humulus* pollen. Biomarker analysis has also been used to identify the pectinolytic microorganisms responsible for the detachment of hemp fibers. For example, DNA- and RNA-based taxonomy has allowed identification of a variety of pectinolytic bacteria and fungi participating in hemp fiber extraction (Ribeiro et al., 2015). Therefore, it would be theoretically possible to record past retting processes by identifying the DNA/RNA sequences of these taxa in past lake sediments. This represents a straightforward evidence of retting, as it is based in the actual occurrence of microorganisms responsible for this biochemical process.

This paper uses sedimentary 16S *ribosomal* RNA genes (16S *rRNA*) from recognized pectinolytic bacteria to demonstrate historical hemp retting during the last five centuries in a Iberian lake where previous *Cannabis* pollen records suggested this possibility but local historical records and oral tradition failed to support it (Rull et al., 2011; Rull & Vegas-Vilarrúbia, 2014). A detailed and high-resolution (subdecadal) hemp pollen record is available for the last centuries in this lake, which was tentatively correlated with the main trends of hemp industry in Spain (Trapote et al., 2018; Rull et al., 2021). In this short communication, the abundance of 16S *rRNA* genes from retting bacteria is analyzed and compared with the available pollen and historical records, detailed analyses on bacterial community dynamics are in progress and will be published further.

## Study area

### The lake

Lake Montcortès is a Pyrenean karstic lake situated at 42° 19’ 50” N - 0° 59’ 41” E and 1027 m elevation (Fig. 1). The regional climate is of montane Mediterranean type and the vegetation is transitional between the submontane and montane belts, which are dominated by oak and pine forests, respectively. The lake is surrounded by a continuous fringe of aquatic vegetation dominated by sedges and reeds (Rull et al., 2011; Mercadé et al., 2013). The sediments of Lake Montcortès are characterized by a long and continuous annually-laminated (varved) sequence encompassing the last ∼3000 years, which has been intensively studied from paleoenvironmental and paleoecological points of view. The record of vegetation and landscape dynamics has been considered unique for the Mediterranean region, in terms of chronological accuracy, resolution and continuity (Rull et al., 2021).

**Figure 1.**
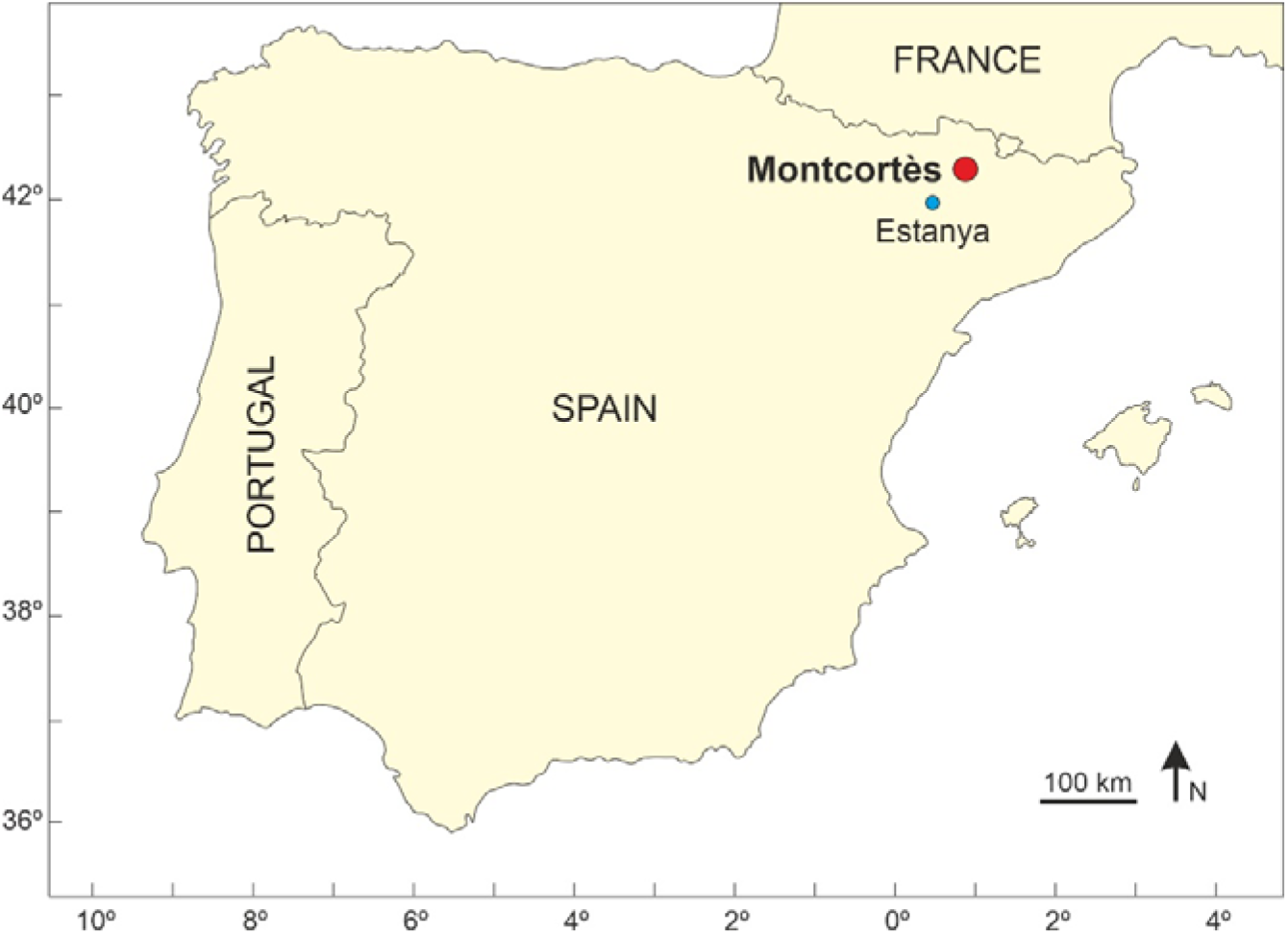
Sketch-map of the Iberian Peninsula with the location of Lake Montcortès (red dot) and Lake Estanya (blue dot).

### Hemp pollen record and historical background

A major feature of this record has been the presence of *Cannabis*/*Humulus* pollen, identified as *Cannabis* on a morphological basis (Rull & Vegas-Vilarrúbia 2014), from Medieval times (∼600 CE) to present. Medieval *Cannabis* records were consistent with local/regional cultivation but the sharp increase experienced during Modern times (16^th^ century) and maintained until the late 19^th^ century was interpreted as evidence of a long phase of hemp retting. These trends coincided with similar Cannabaceae records in the neighbor Lake Estanya (Riera et al., 2004, 2006), and were consistent with the general development of hemp industry across the whole country (Spain), as documented in historical records. However, historical evidence of retting practices in Lake Montcortès and its surroundings was lacking in written documents and oral tradition (Marugan & Rapalino, 2005).

The most detailed hemp pollen record from Lake Montcortès was obtained in core MONT-0713-G05 and is displayed in Fig. 2. Percentages were below 10% during the first half of the 16^th^ century and suddenly increased beyond the purported retting boundary (15-25%) shortly after 1550 CE to remain in this situation until the end of the 17^th^ century. According to Sanz (1995), hemp cultivation in most of the Iberian Peninsula (IP) during the 16^th^ and 17^th^ centuries was primarily oriented to supply fiber for domestic needs and for local trading. However, the significant growth of the Spanish royal navy experienced after the Columbian discovery of America (1492 CE) notably increased the demand for hemp fiber, mainly for sails and ropes, which coincided with the 16^th^-17^th^ century hemp pollen increase in the Montcortès record. A second acceleration in this curve took place at the beginning of the 18^th^ century, coinciding with the explicit mandate of cultivating hemp in every land suitable for this purpose and selling the harvests to the crown for the growth of the Royal Navy (Sanz, 1995).

**Figure 2.**
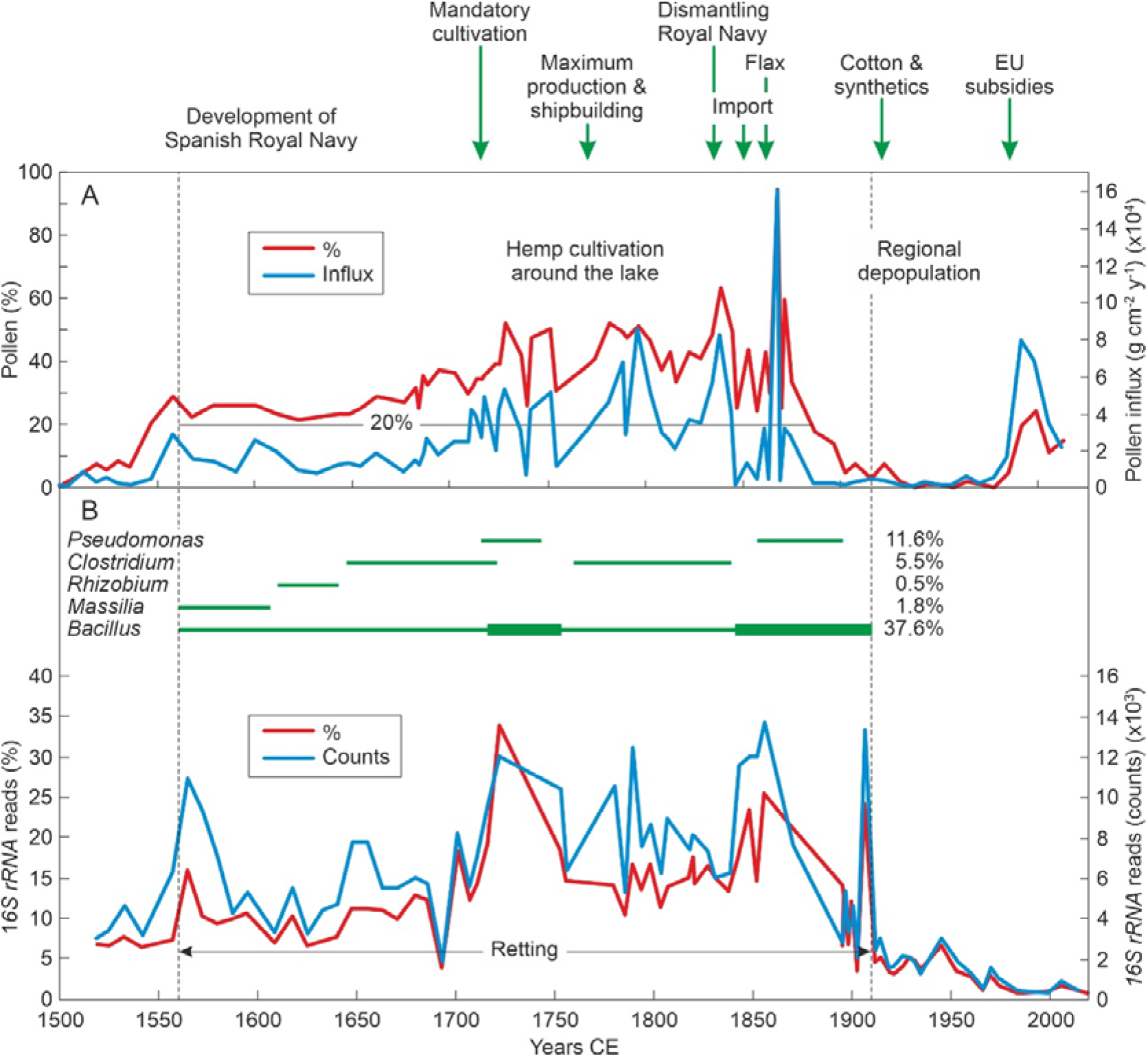
Comparison of hemp pollen and phylogenetic bacterial records in Lake Montcortès for the last five centuries, in relation to the main developments of the Spanish hemp industry. A) Hemp pollen record (core MONT-0713-G05) in percentage and influx units. Note that the threshold pollen percentage within the retting phase is approximately 20%. Redrawn and modified from Trapote et al. (2018) and Rull et al. (2021). B) Results of 16S *rRNA* analysis performed in this paper (core MONT-0119-11) expressed as the total percentage and counts of phylogenetic material from retting bacteria. The main taxa participating in the retting process are represented as horizontal green lines indicating presence/absence phases (maximum percentages for each taxon are given in the right side). Raw data for this figure is provided as Supplementary Material (raw data).

The Montcortès hemp curve remained significantly above the pollen-inferred retting boundaries until late-19^th^ century, a period characterized by local and regional hemp cultivation around the lake (Madoz 1845-50; Ferrer, 2017). This was the period of maximum hemp production and shipbuilding in Spain (1750-1775 CE), which has been attributed to a phase of general economic prosperity and intensification of commerce with America (Andreu, 1981; Delgado, 1994). A drastic shift occurred in 1834 CE with the decadence of the Spanish Empire and the dismantling of the Royal Navy. This event was clearly reflected in the Montcortès hemp curve, which underwent a sharp decline shortly before 1850 CE. Imports from other countries contributed to a significant drop in hemp cultivation, which was substituted by other products, including alternative fiber products such as flax, since 1860 CE (Casassas, 1985). The coincidence of the Montcortès hemp pollen trends with the main events of the hemp industry, at the country level, suggests that Lake Montcortès could have been an important regional retting place for almost four centuries.

Besides the occurrence of a percentage and influx peak by ∼1870 CE - which was provisionally attributed to possible sediment disturbances caused by a phase of frequent heavy rainfall events (Trapote et al., 2018) – the second half of the 19^th^ century and most of the 20^th^ century were characterized, in the Montcortès record, by the absence of hemp pollen, suggesting the cessation of any activities linked to the hemp industry. This was likely caused by the abandonment of the lake catchment, which has been linked to the vast Pallars depopulation caused by the crisis of subsistence economy (1870-1910 CE) and the further massive emigration to industrial cities (1960-1980 CE), which resulted in a 60% reduction of the Pallars population (Farràs, 2005).

Hemp pollen reappeared in Lake Montcortès sediments in the 1990s but neither cultivation nor retting of this plant were active in the lake surroundings, which suggests the existence of extra-local regional sources. As demonstrated in the study of modern pollen deposition, *Cannabis* is currently present throughout the year in significant percentages (Rull et al., 2017), even in the absence of local cultivation and retting practices, which has been attributed to the high dispersion power of this anemophilous pollen (Cabezudo et al., 1997; Munuera et al., 2006). Therefore, although the exact source of regional hemp pollen in modern sediments could not be located, it is assumed that it exists somewhere. The same could be valid for the *Cannabis* pollen increase of the last decades, as documented in the pollen record, which coincides with a general country-level expansion of hemp cultivation, favored by European Union subsidies to this activity (Karus & Kaup 2002; Gorchs & Lloveras 2003).

## Methods

This work compares the above hemp pollen record, obtained in core MONT-0713-G05 (Fig. 2), with the sedimentary 16S *rRNA* genes record of well-known retting bacteria from a newly retrieved core (MON-0119-11). These parallel cores were recovered a few meters apart in the same location (42° 19’ 54.4” N - 0° 59’ 42.1” E; 28.2 m of water depth) and were stratigraphically analogous. The age-depth model of core MON-0119-11 (Fig. 3) was built by correlation with core MONT-0713-G05 using key marker horizons and characteristic varve sequences.

**Figure 3.**
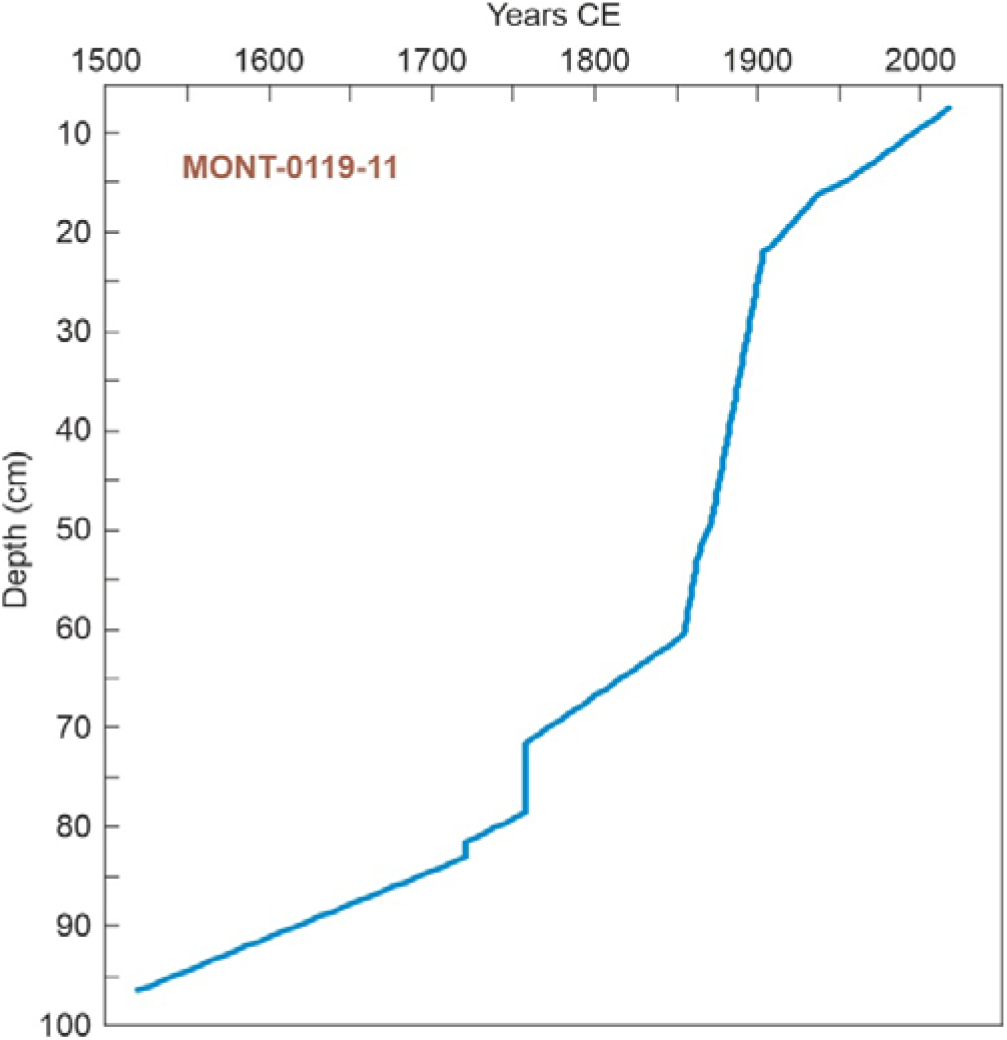
Age-depth model for core MON-0119-11.

The pectinolytic bacteria analyzed in this work were those considered to be the most relevant in the hemp retting process, which were previously characterized using 16S *rRNA* genes (Ribeiro et al., 2015), namely: *Bacillus/Paenibacillus, Clostridium, Escherichia, Pseudomonas, Massilia, Rhizobium, Rhodobacter* and *Methylobacterium*. Sequences taxonomically ascribed to the same taxa were grouped together. Results are expressed as read counts and as percentages with respect to the total read counts (>0.1% abundance and prevalence). DNA was extracted from samples collected at 1-cm intervals from the sediment core using the FastDNA Spin Kit for Soil (MP Biomedical) according to the manufacturer’s instructions. High-throughput multiplexed 16S *rRNA* gene sequencing with the Illumina MiSeq System (2×250 PE) was carried out using primer pair 515f/806r (Caporaso et al., 2011) targeting the V4 region of the 16S *rRNA* gene complemented with Illumina adapters and sample specific barcodes. Details on the analysis of sequence datasets are described in the Supplementary Material (methods).

## Results and interpretation

The total abundance of 16S *rRNA* genes from retting bacteria increased in the mid-16^th^ century and stabilized until the early 18^th^ century, when it experienced an abrupt increase and a further decline (mid-18^th^ century) followed by another increase during the second half of the 19^th^ century and a sharp decline in the early 20^th^ century, with no further recovery (Fig. 2). These trends indicate that hemp retting was ongoing in Lake Montcortès between mid-16^th^ and early-20^th^ centuries, with a maximum during the first half of the 18^th^ century and the second, less pronounced increase in the second half of the 19^th^ century. Therefore, previous inferences about the occurrence of hemp retting based on indirect evidence – i.e., pollen analysis (Rull & Vegas-Vilarrúbia, 2014; Rull et al., 2011, 2021; Trapote et al., 2018) – are confirmed by direct evidence of this activity. In addition, trends in bacterial abundance are consistent with *Cannabis* pollen abundances, which were above 20% during the whole retting phase (except for the last decades, as further discussed), with values between 25% and 60% during the interval of higher retting intensity (18^th^ and 19^th^ centuries). This is in agreement with the 15-25% hemp pollen threshold used by several authors to infer local retting practices on the basis of pollen abundances (Lavrieux et al., 2013; McPartland et al., 2018).

The good match between retting activity, as recorded by bacterial 16S *rRNA* genes, and hemp pollen abundance also reinforce the previously-established general relationships between the hemp pollen record and the historical development of the hemp industry in Spain (Fig. 2). For example, hemp retting in Lake Montcortès initiated during the development of the Royal Navy and peaked just after the mandatory nature of hemp cultivation across the whole country (Sanz, 1995). Also noteworthy is the coincidence of the retting cessation with the dramatic depopulation of the Pallars region, in the 20^th^ century. These observations suggest that Lake Montcortès could have been an important retting center for the hemp produced at regional level for more than four centuries, especially during 18^th^ and 19^th^ centuries, a hypothesis that should be tested with further studies.

Some minor differences between pollen and bacterial dynamics curves are worth to mentioned. One is that the final drop occurred several decades before in the pollen record (late 19^th^ century) than in the bacterial one (early 20^th^ century), which may suggest the continuity of retting practices in the absence of hemp. As mentioned above, hemp cultivation underwent a significant decline after the dismantling of the Spanish Royal Navy and was replaced by other fiber sources such as flax, since 1860 CE (Casassas, 1985). Therefore, it is possible that hemp retting was replaced by flax retting at the end of the Montcortès retting period. Further paleoecological and historical studies should consider this possibility. Another difference is that the pollen increase recorded during the last decades (1980-90 CE onward) did not occur in the bacterial curve, indicating that this pollen should have originated in extra-local sources and reached the lake by wind dispersal (Rull et al., 2017). The above-mentioned increase in hemp production fostered by EU subsides (Karus & Kaup 2002; Gorchs & Lloveras 2003) may be involved in this phenomenon but the exact source of hemp pollen in this case remains unknown.

The trends in total retting bacteria were largely due to changes in *Bacillus*/*Paenibacillus*, which was the most abundant taxon, by large, accounting for up to 38% of the total bacterial community, and was present throughout the entire retting phase (Fig. 2). Other taxa were less abundant and continuous, showing discontinuous presence/absence phases that conform a clear long-term successional pattern. The retting phase began with the succession *Massilia-Rhizobium-Clostridium-Pseudomonas* until the 18^th^ century peak. After this peak, the succession restarted but that time only the Clostridium-Pseudomonas stages occurred before reaching the second, less intense retting peak. This may suggest that the less abundant taxa (*Massilia* and *Rhizobium*) were only present at the beginning of the retting process and, once this process was ongoing, these initial stages were skipped. The other retting taxa analyzed (*Escherichia, Methylobacterium* and *Rhodobacter*) occurred only on scattered samples, in very low percentages, without evident presence/absence patterns.

Hemp retting is known to be a major anthropogenic disturbance for the water quality and the ecological dynamics of the affected waterbodies, especially in relation to eutrophication (Iwańska et al., 2022). In Lake Montcortès, hemp retting has been related with eutrophication processes, as documented in algal (desmids, diatoms) and macrophytic (*Typha*) records (Scussolini et al., 2011; Trapote et al., 2018). Detailed analyses of RNA-based bacterial community dynamics are in progress to provide more information about the potential influence of hemp retting on the ecological functioning of the lake.

## Conclusions and further research

The analysis of sedimentary 16S *rRNA* genes from bacteria involved in the process of hemp retting demonstrated that this process occured in Lake Montcortès between mid-16^th^ and early 20^th^ centuries, with a maximum in the first half of the 18^th^ century. These results follow similar patterns of previous hemp pollen records and are closely related to the general trends of the Spanish hemp industry during the last five centuries. Hemp pollen remained above 20% during the whole retting phases, suggesting that this is the pollen threshold to infer hemp retting in this lake, which falls within the range of previous studies from other areas (15-25%). The results of this study can be useful to interpret paleoecological sequences from other similar lakes in the way toward a more regional view on the introduction, spreading, uses and associated cultural connotations of *Cannabis* in the Iberian Peninsula within European and Mediterranean contexts (Rull, 2022).

## Acknowledgments

This works has been funded by the Spanish Ministry of Economy and Competitiveness, projects MONT-500 (CGL2012-33665) and MEROMONT (CGL2017-85682-R).

## Supplementary Material

Detailed methods of phylogenetic analysis and the raw data obtained are provided as supplementary word and excel files.

## Author contributions

VR: conceptualization, methodology, formal analysis, investigation, data curation, writing, visualization. OSS: methodology, formal analysis, data curation, review & editing. ASM: methodology, formal analysis, data curation. CMB: methodology, formal analysis, data curation, review & editing. TVV: investigation, resources, review & editing, funding acquisition.

